# TimTrack: A drift-free algorithm for estimating geometric muscle features from ultrasound images

**DOI:** 10.1101/2020.08.23.263574

**Authors:** Tim J. van der Zee, Arthur D. Kuo

## Abstract

Ultrasound imaging is valuable for non-invasively estimating fascicle lengths and other features of pennate muscle, especially when performed computationally. Effective analysis techniques to date typically use optic flow to track displacements from image sequences, but are sensitive to integration drift for longer sequences. We here present an alternative algorithm that objectively estimates geometric features of pennate muscle from ultrasound images, without drift sensitivity. The algorithm identifies aponeuroses and estimates fascicle angles to derive fascicle lengths. Length estimates of human vastus lateralis and lateral gastrocnemius in healthy subjects (N = 9 and N = 1 respectively) compared well (root-mean-square error, RMSE < 0.80 cm) to manual estimates by independent observers (n = 3). The coefficient of multiple correlation (CMC) with manual estimates of fascicle length was comparable to previously reported for state-of-the-art optic flow algorithm (0.93-0.99), suggesting similar accuracy. The algorithm requires minimal manual intervention and can optionally extrapolate fascicle lengths that extend beyond the image frame. It facilitates automated analysis of ultrasound images without drift.

## Introduction

Ultrasonography, or ultrasound, can be used to non-invasively estimate features of pennate muscle geometry, including muscle thickness [e.g. 1], pennation angle [e.g. 2] and fascicle length [e.g. 3]. These features are of particular interest in biomechanics, because they influence both mechanical [e.g. 4–6] and energetic aspects of muscle force production [e.g. 7–9]. Whereas ultrasound methods was originally performed manually(semi-) automated methods have recently become more prevalent [for a recent review, see 10]. Unfortunately, existing (semi-)automated methods that estimate fascicle length are prone to drift due to using optic flow, still rely on considerable user interaction, and/or are not freely available.

One of the challenges with automated ultrasound analysis is image speckle, a type of interference that degrades image quality and hampers tracking of single fascicles [10]. Sensitivity to image speckles can be reduced by using optic flow, which is one of the primary (semi-)automated methods. Optic flow uses the image velocity field to indicate bulk changes that are relatively insensitive to speckle, and thus estimates fascicle length changes between succeeding image frames [11–13]. One shortcoming is that accuracy requires succeeding images to be similar, such that faster muscle contraction requires high image capture rates [10]. Another drawback is that optic flow techniques accumulate the frame-by-frame changes over time, which also accumulates errors over long sequences, referred to as integration drift. Optic flow algorithms are therefore also history dependent, and usually need an initial, manually-tracked image that may introduce subjectivity from human operators. A final, manually-tracked image at the end of an image sequence may also be used to reduce drift, but again with some subjectivity. Optic flow techniques are therefore most effective for relatively short image sequences captured at high rates, and typically require significant user intervention.

Another approach is automated feature detection within a single image, rather than optic flow across multiple images. Detected features may include muscle thickness [14], fascicle orientation [15–18], pennation angle [19] and fascicle length [20]. To reduce influence of speckles, feature detection algorithms typically rely on image filtering procedures. Filtered images may be used to identify individual fascicle objects (Marzilger et al., 2018) and features such as fascicle orientation, or to obtain aggregate feature estimates using line detection procedures (Ryan et al., 2019). Although such detection is history independent, most feature detection algorithms still rely on some degree of manual intervention, for example to detect the aponeuroses [e.g. 17, 18], which helps to define fascicle lengths. However, others have used feature detection techniques to also automate aponeurosis detection (Zhou et al., 2015). It is thus possible to automate most detection steps, yielding a more objective, repeatable, and convenient means of estimating muscle fascicle features without drift.

Here we present an algorithm called TimTrack that automatically estimates geometric features of pennate muscle. The algorithm uses a combination of feature detection methods, to specifically determine muscle thickness, muscle fascicle orientation, pennation angle, and fascicle length without drift. The algorithm estimates fascicle orientations using a line detection procedure [21] as in other algorithms [15, e.g. 18, 20], and automatically detects aponeuroses to estimate muscle thickness. Geometric calculations combine these data to estimate fascicle lengths and pennation angles. With appropriate geometric assumptions, the algorithm can also extrapolate to estimate fascicle lengths that extend beyond an ultrasound image. We here show that TimTrack’s estimates are comparable to those from both manual observers and a commonly used optic flow algorithm.

## Methods

The developed TimTrack (feature detection) algorithm estimated geometric muscle features from ultrasound images. First, the algorithm employs a filter to highlight and detect aponeuroses in the images. It then uses a technique called the Hough transform to estimate the overall fascicle orientation. These data, together with geometric calculations, yield estimates of muscle thickness, pennation angle, and muscle fascicle length. We tested the algorithm on ultrasound images obtained from vastus lateralis and lateral gastrocnemius muscle in healthy human subjects and compared its estimates to manual estimates from independent observers (N=3).

## Algorithm

There are four main steps to the algorithm (see Fig 1). Prior to its application, there is a manual (0.) preparation step in which the user specifies two image regions of interest where the aponeuroses are to be detected. The subsequent steps are automated: (1.) Highlight line-like structures for fascicles (thin lines) and aponeuroses (thick lines). (2.) Detect aponeuroses and their inner edges. (3.) Determine fascicle angles. (4.) Estimate muscle thickness *T*_muscle_, pennation angle *φ* and fascicle length *L*_fas_. There are also two optional steps that may be performed, depending on the application: (5.) Extrapolate geometry beyond the image frame for longer fascicles. (6.) Time-interpolate aponeuroses for image sequences with missing data (e.g. from occlusion). These steps are explained in greater detail next.

**Fig 1:**
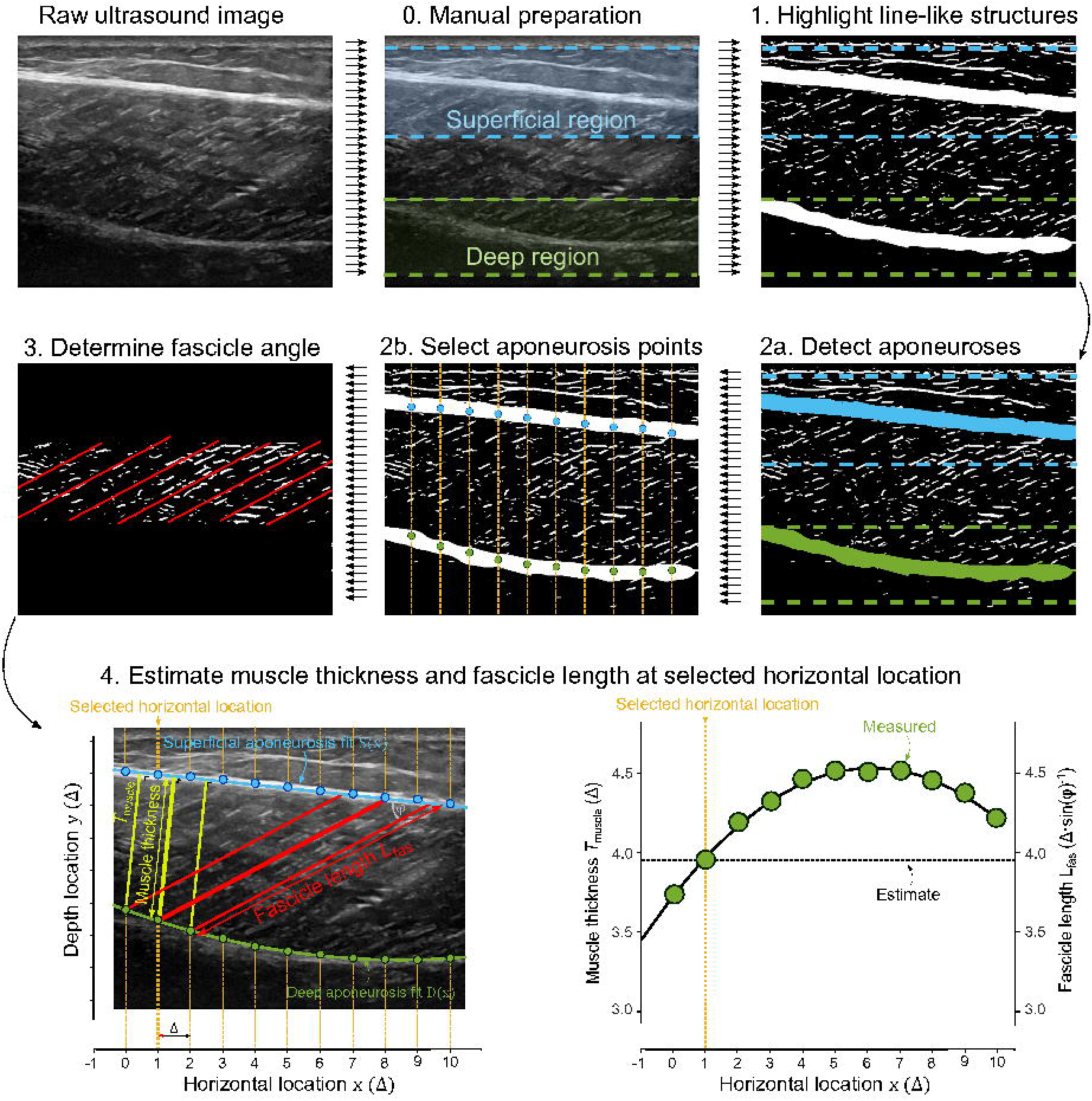
Summary of steps in processing of ultrasound images. Raw images undergo manual preparation to select regions, and image filtering to highlight line-like structures. Aponeuroses are detected from the filtered images and specific points are selected on the aponeuroses (green and blue dots) at specified horizontal locations (vertical yellow lines). The region between aponeuroses is used to determine fascicle angle using the Hough transform. Pennation angle *φ* is estimated from fits through the identified superficial aponeurosis points and the fascicle angle. Muscle thickness *T*_muscle_ is defined as the perpendicular distance from the deep aponeurosis to the (fitted) superficial aponeurosis. Fascicle length *L*_fas_ is calculated from pennation angle *φ* and muscle thickness *T*_muscle_ using trigonometry. Muscle thickness *T*_muscle_ and therefore fascicle length *L*_fas_ can be evaluated at specific horizontal location *x*.

### 0. Preparation

Before applying the algorithm, the user selects two regions of interest where the superficial and deep aponeuroses are to be detected. This assumes an image where the superficial aponeurosis is near the top, and the deep aponeurosis is near the bottom. The regions are specified by a relative depth range for each aponeurosis (*D*_superficial_ and *D*_deep_, see Supporting Information). In practice, these regions need normally be specified only once for an entire image sequence.

### 1. Highlight line-like structures

Line-like structures are detected within the image using a medical imaging technique termed the Frangi-type vessel enhancement filter [22]. Such filtering was originally developed for enhancing blood vessels in imaging and has later been applied to enhancing line-like muscle fascicles [16, 18] and aponeuroses [14] on separate occasions. We here apply the filter to both muscle fascicle and aponeurosis detection. We adapted an open-source implementation of this filter (Dirk-Jan Kroon, Matlab Central, 2021) to reduce edge effects, following a user suggestion to “adding the ‘replicate’ argument to the imfilter call in Hessian2D.m” (user Phillip, Matlab Central). The adapted filter is applied twice, using different line thickness settings for aponeuroses (thick) and muscle fascicles (thin) (σ_fas_ and σ_apo,H_, see Supporting Information), to yield two filtered images. The filtered fascicle image (step 1.1a, Fig 2) is thresholded (step 1.2a, Fig 2, *T*_fas_, see Supporting Information), and the filtered aponeurosis image (step 1.1b, Fig 2) is masked (step 1.2b, Fig 2). The mask is created by thresholding the original image (step 1.1c, Fig 2, *T*_apo_, see Supporting Information), and applied by multiplying against the aponeurosis image, to reduce boundary effects of the vessel enhancement filter. Next, a second mask is applied to retain only the user-defined depth region for each aponeurosis and the remaining white pixels in each depth region (i.e. superficial and deep) are smoothed using a 2D Gaussian kernel (*imgaussfilt*, MATLAB, step 1.3, Fig 2, σ_apo,k_, see Supporting Information). Finally, the filtered aponeurosis image is thresholded (step 1.4, Fig 2) and filtered so that only the longest two objects in each depth region remain. An object is defined as a collection of connected white pixels with continuous adjacency. If the length ratio between the smallest and longest of these two objects is less than a pre-set parameter value (*L*_ratio,max_, see Supporting Information), the longest object is coined the aponeurosis object (*bwpropfilt*, MATLAB, using the *Major Axis Length property*). If the length ratio is larger than the parameter value, the objects are used to mask the grayscale (i.e. non-thresholded) image. The object resulting in the masked grayscale image with the highest mean pixel intensity is coined the aponeurosis object (*bwpropfilt*, MATLAB, using the *Mean Intensity property*). The aponeurosis objects are used for aponeurosis detection and subtracted from the fascicle image (step 1.5, Fig 2), which is subsequently used for the Hough transform.

**Fig 2.**
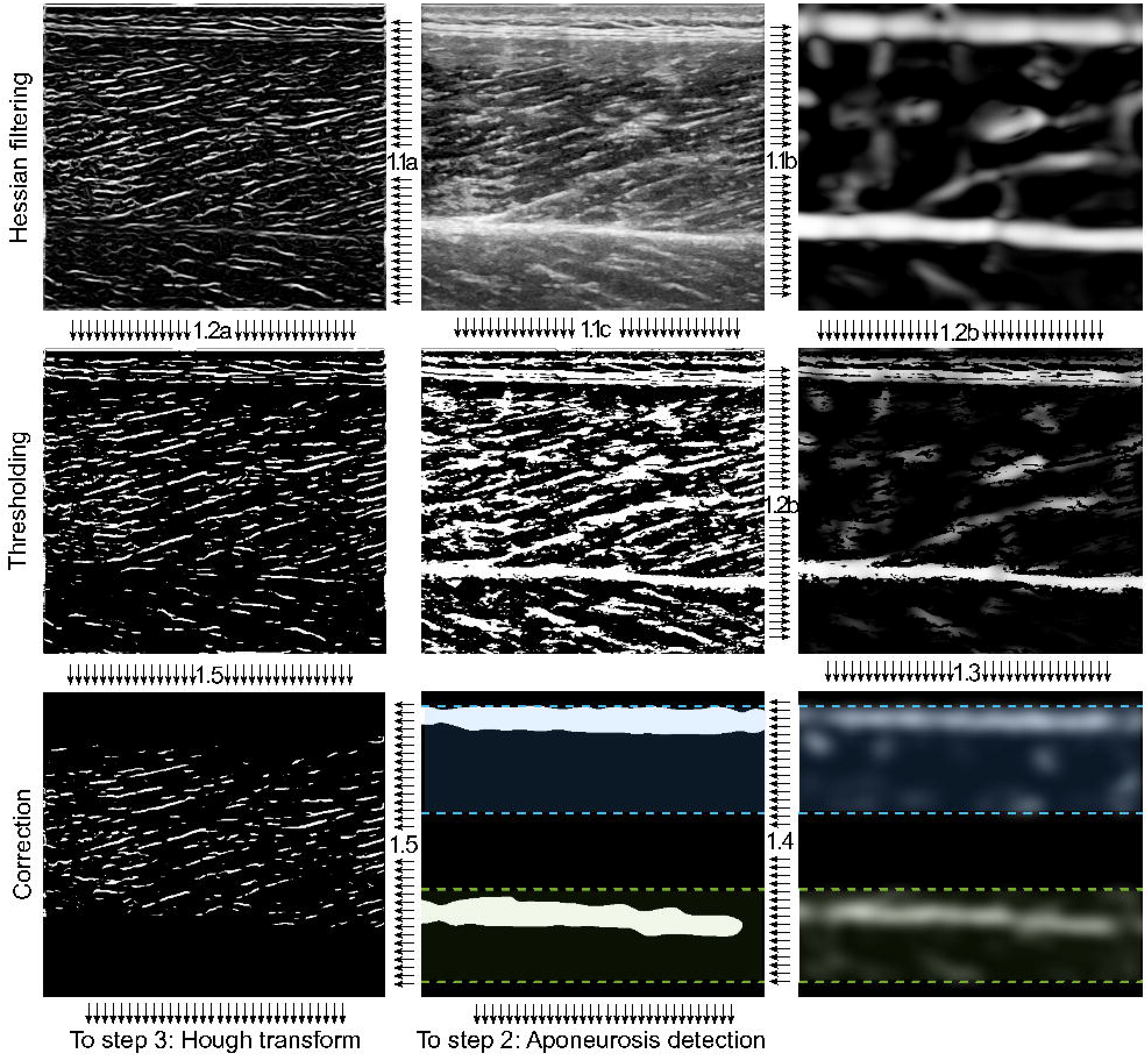
Image filtering steps for highlighting line-like structures. Filtering thin lines for fascicle detection (step 1.1a) and thick lines for aponeurosis detection (step 1.1b). Fascicle filtering includes thresholding (step 1.2a) and correction for aponeurosis pixels (step 1.5). Aponeurosis filtering includes a combination of thresholding with Hessian filtering (step 1.2b), Gaussian smoothing (step 1.3) and second thresholding (step 1.4). The results of the fascicle and aponeurosis filtering process are inputs for the Hough transform (step 3) and aponeurosis detection respectively (step 2).

### 2. Detect aponeuroses and their inner edges

In the second step, the inner edges of the superficial and deep aponeurosis objects are determined, and used to define sampled aponeurosis points. The inner edges are defined as the white pixels on the aponeurosis objects closest to the interior of the image. The edges are first trimmed to compensate for the width added by the 2D Gaussian kernel used in step 1.3. The superficial and deep aponeurosis points are defined as these corrected inner edges, evaluated at a predefined number of equidistantly spaced horizontal locations (*n*_apox_, see Supporting Information) across the image width (solid dots in Fig 1), with specified margin from the image boundary (*x*_margin_, see Supporting Information). The latter allows for cropping from each side to reduce edge effects. The superficial and deep aponeurosis are linearly interpolated and extrapolated using superficial and deep aponeurosis fits *y* = *S*(*x*) and *y* = *D*(*x*) respectively (solid lines in Fig 1). Bounds may be placed on the maximal value of the fits’ derivative (specified by *β*_max_ and *γ*_max_, see Supporting Information) to avoid physiologically implausible fits, using constrained optimization (MATLABs *fmincon*). The aponeurosis fits can be used to evaluate the (depth locations *y* of the) aponeuroses at any horizontal location *x* (inset Fig 1), even if it is outside the image frame (i.e. extrapolation). The superficial aponeurosis fit *S*(*x*) is always linear, while the deep aponeurosis fit *D*(*x*) is a polynomial with predefined order (*o*_deep_, see Supporting Information). The former simplifies quantifying superficial aponeurosis angle and pennation angle, while the latter allows for incorporating the deep aponeurosis curvature into the muscle thickness estimate.

### 3. Determine fascicle angles

In the third step, fascicle angle is determined by applying the Hough transform to the filtered fascicle image (see Fig 3). First, the fascicle region-of-interest is defined as the region interior to the aponeuroses, defined by the average vertical depth of the aponeurosis points. Before the Hough transform is applied, an ellipse is cut out of the (rectangular) fascicle region-of-interest (step 3.1, Fig 3). This is to reduce bias towards diagonal lines, which typically occurs when applying the Hough transform to rectangular images [e.g. 23]. The ellipse width *w*_ellipse_ is a fraction of the fascicle image width (*w*_ratio_, see Supporting Information). The horizontal and vertical coordinates of the ellipse center are equal to half the ellipse width *w*_ellipse_ and the fascicle image height respectively. The Hough transform is applied to the ellipsoid fascicle image (*hough*, MATLAB), resulting in a 2D accumulator matrix (step 3.2, Fig 3). The 2D accumulator matrix represents frequency counts of lines that are parameterized by angle *θ* and distance *ρ*, using a predefined angle resolution (*θ*_res_, see Supporting Information), angle range (*θ*_range_, see Supporting Information) and distance resolution of 1 pixel. Angle *θ* is the angle of a vector running from the origin to the line along a vector perpendicular to the line, distance *ρ* is its length. The Hough angle *θ* sets the angle of the line itself, which we call *γ*:

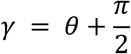

**Fig 3.**
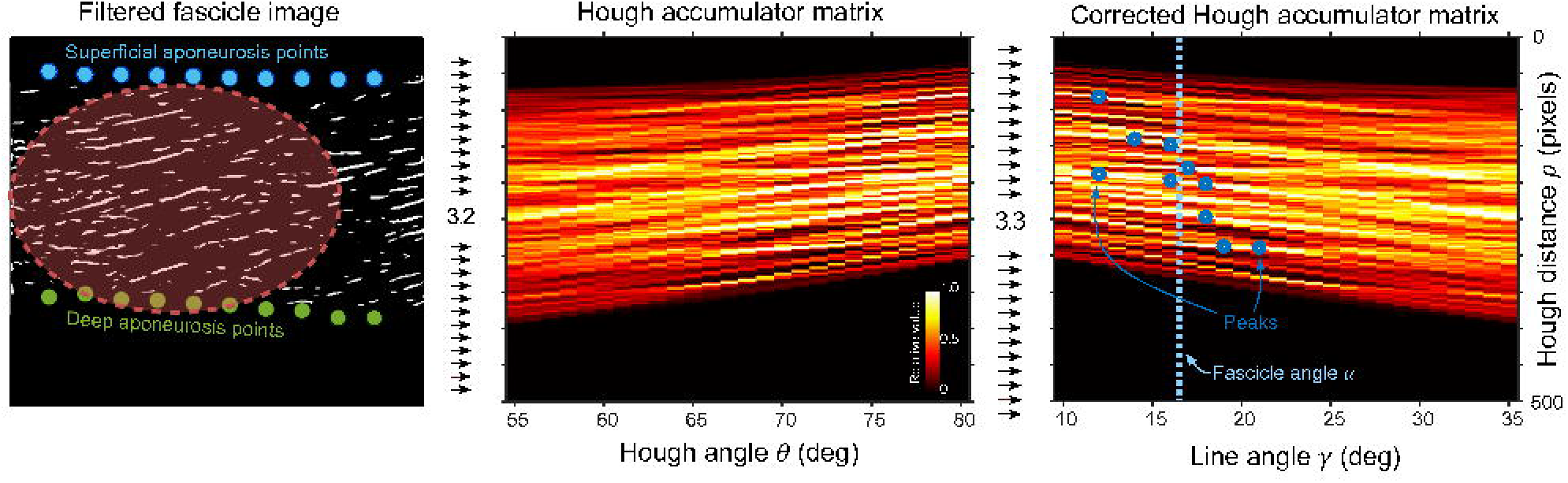
Use of Hough transform to determine fascicle angle. Fascicle depth region is determined from superficial and deep aponeurosis points and an ellipse is cut out (step 3.1). The Hough transform is performed on the ellipsoid image, resulting in a 2D accumulator matrix (step 3.2). The accumulator value is corrected for the corresponding radius of the ellipse and the peak values are determined (step 3.3). Fascicle angle α is defined as the median of the line angles corresponding to the peaks.

Next, the 2D accumulator matrix is corrected for the effect of angle *δ* on the relative ellipse radius *r*_ellipse,rel_ (step 3.3, Fig 3). The peaks in the corrected 2D accumulator matrix are determined (*houghpeaks*, MATLAB) and the angles *γ* belonging to the highest peaks are selected, using a pre-set number of included peaks (*K*, see Supporting Information). Fascicle angle *α* is defined as the median of the selected angles *γ*_selected_ (step 3.4, Fig 3). We chose to use the median instead of the mean for better robustness to outliers.

### 4. Calculate muscle thickness, pennation angle and fascicle length

In the fourth step, pennation angle *φ* and muscle fascicle length *L*_fas_ are determined from aponeurosis fits and fascicle angle *α*. Pennation angle *φ* is defined as the difference between superficial aponeurosis angle *β* and the fascicle angle *α*,

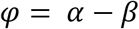

Superficial aponeurosis angle *β* is defined as the angle of the (linear) superficial aponeurosis fit *S*(*x*). Muscle thickness *T*_muscle_ is defined as the perpendicular distance from the (fitted) deep aponeurosis to the (fitted) superficial aponeurosis, and can be evaluated at any horizontal location *x* (see Fig 1):

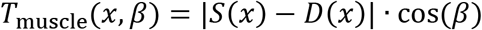

Fascicle length *L*_fas_ is calculated from pennation angle *φ* and muscle thickness *T*_muscle_ using trigonometry (see Fig 1),

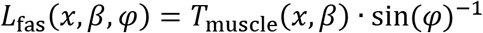

### 5. Extrapolate geometry beyond the image frame for longer fascicles (optional)

If the muscle fascicles extend beyond the width of the ultrasound image, the algorithm can linearly extrapolate the aponeuroses and fascicles (see Fig 4). In the program, the user may switch from ‘regular mode’ (without extrapolation) to ‘extrapolation mode’. In extrapolation mode, the selected horizontal location *x* for each image is chosen such that the extrapolation is spread between the left and the right sides of the image. This is accomplished by requiring the fascicle of interest to go through the midpoint *M* of the image (see Fig 4). The selected horizontal location *x* is then defined as the horizontal coordinate of the intersection between the extrapolated deep aponeurosis and the extrapolated fascicle of interest. For our data set, we used ‘regular mode’ and chose to set the horizontal location *x* to the left image border to facilitate comparison with manual estimates of muscle thickness at this location. The ‘extrapolation mode’ is demonstrated here with the vastus lateralis, which had relatively long fascicles extending beyond the frame of our images.

**Fig 4.**
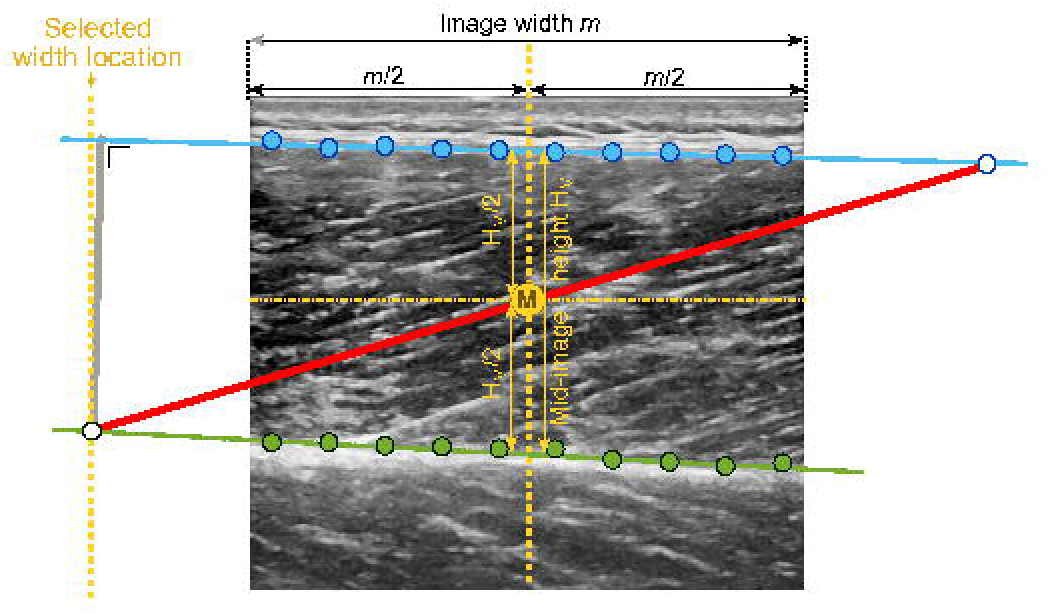
Extrapolation of muscle fascicle beyond image frame. The fascicle of interest (thick red line) goes through the midpoint *M*, and intersects with aponeuroses at locations outside of the image frame. The horizontal coordinate of the midpoint *M* is halfway the image width *m*. The vertical coordinate of the midpoint *M* is halfway between the deep and superficial aponeuroses. The selected width is determined by finding the intersection between the extrapolated fascicle of interested and the extrapolated deep aponeurosis.

### 6. Time-interpolate aponeuroses for image sequences with missing data (optional)

Image quality may deteriorate with excessive muscle contraction velocity (image blur) or when part of the ultrasound probe loses contact with the skin (image occlusion). In these cases, aponeuroses may only be partially visible, resulting in missing aponeurosis points. When analyzing image sequences, these missing data points may be replaced using time-interpolation. To reduce the effect of misidentified aponeurosis points, our custom (post-hoc) interpolation procedure only includes points that are less than a predefined number of standard deviations away from the mean (see Table 1).

## Experimental testing of algorithm

We employed the algorithm on ultrasound images obtained from human vastus lateralis and lateral gastrocnemius muscle. In a first experiment, ultrasound images of the lateral gastrocnemius muscle in one human subject were recorded while (1) actively going through the range-of-motion and (2) jumping, both at an image capture rate of 25 Hz. In a second experiment, ultrasound images of vastus lateralis muscle in human subjects (6 male, 3 female, leg length = 87.4 ± 4.9 cm, body mass = 70.6 ± 13.1 kg, mean ± s.d.) were recorded during isometric torque production, at an image capture rate of 30 Hz. The resulting estimates were low-pass filtered to reduce the effects of random noise, using cut-off frequencies of 1 Hz for gastrocnemius range-of-motion and vastus lateralis torque production, and 10 Hz for gastrocnemius jumping. The algorithm’s estimates were compared to manual estimates from three independent observers, quantified with the root-mean-square error (RMSE). To assess the algorithms (dimensionless) accuracy relative to current state-of-the-art algorithms, the coefficient of multiple correlations (CMC) [24] between the algorithms estimates and manual observer estimates was computed. Fascicle length and angle (and corresponding accuracy measures) were also estimated with a commonly used optic flow algorithm (i.e. UltraTrack) [12]. This was only done for gastrocnemius images and not the vastus lateralis images, which required extrapolation beyond the image frame. To assess the variability within manual observer estimates, the difference between manual estimates of different observers was quantified using the root-mean-square difference (RMSD). The intraclass correlation coefficient (ICC) between manual observer estimates was computed to assess the difference in manual observer estimates relative to previously reported values. Linear regressions were performed for muscle thickness, pennation angle and fascicle length with mean of manual estimate as independent variable and algorithm estimate as dependent variable. This was done to assess potential systematic error, which would be absent if the linear regression coefficient equals unity (1). In the remaining, we refer to the current algorithm as TimTrack.

## Results

The TimTrack algorithm captured changes in gastrocnemius fascicle angle and fascicle length relatively well during range-of-motion movement. The algorithm performed well in comparison to manual tracking by human observers, and in comparison with the well-known UltraTrack algorithm (see Fig 5). There were some cases where TimTrack appeared to perform well compared to optic flow, for example at the extremes of motion when there were relatively fast transient accelerations, and relatively large fascicle angles and small fascicle lengths (from comparison with mean of manual estimates). In addition, TimTrack appeared to be less sensitive drift in its estimates of fascicle length towards the end of the range-of-motion trial.

**Fig 5.**
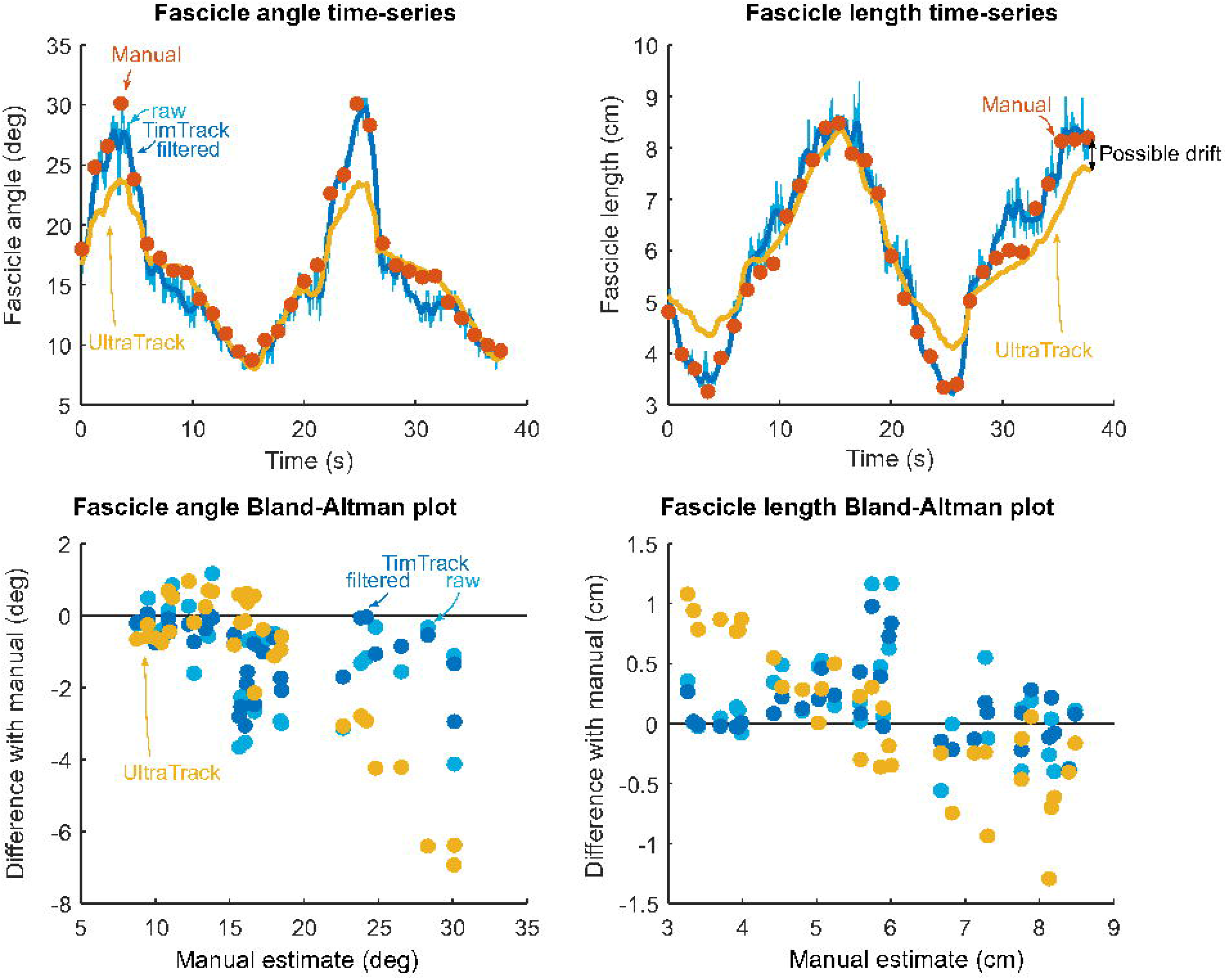
Gastrocnemius fascicle angle and fascicle length estimates of TimTrack algorithm, manual observers and UltraTrack during range-of-motion movement. Upper row: fascicle angle and -length changed considerably during range-of-motion movement, captured by manual observers (red dots) current algorithm (blue line) and UltraTrack (yellow line). UltraTrack underestimated fascicle length near the end of the trial, potentially due to integration drift. Lower row: Bland-Altman plot indicates errors of current algorithm’s raw estimates (light blue dots), filtered estimates (dark blue dots) and UltraTrack’s estimates (yellow dots) as a function of manual estimates.

Manual estimates of muscle thickness, pennation angle and fascicle were quite consistent across human observers (Table 2). TimTrack algorithm estimates were generally comparable to manual estimates, as shown for typical example of each trial (see Fig 6). Gastrocnemius fascicle angle changed more than 3-fold (from about 10° to about 30°) in both range of motion (Fig 6A) and jumping trials (Fig 6B), while changes in superficial aponeurosis angle were less pronounced (Fig 6D–E). Mostly due to the changes in fascicle angle, pennation angle changed considerably during both trials (from about 15° to about 35°, Fig 6G–H). Muscle thickness had some subtle changes during both trials, that appear to be present in both manual and algorithm estimates (Fig 6J–K). Due to considerable pennation angle changes (Fig 6G–H) at near constant muscle thickness (Fig 6J–K), muscle fascicle length changed more than 2-fold in both trials (from about 11 cm to about 4 cm, Fig 6M–N). Vastus lateralis fascicle angle (Fig 6C), superficial aponeurosis angle (Fig 6F) and pennation angle (Fig 6I) increased during the isometric torque production experiment. VL muscle thickness initially increased and later decreased (Fig 6L). Fascicle length decreased over time (see Fig 6O), with a larger decrease at the start than at then end of the trial. All effects were captured by both the TimTrack algorithm (blue dots in Fig 6) and the manual observers (mean as red dot, range as red vertical lines in Fig 6), indicating that TimTrack was qualitatively adequate.

**Table 2:** Differences between manual observer estimates of muscle thickness, pennation angle and fascicle length in ROM, jumping and isometric trials.

|  | Muscle thickness |  |  | Pennation angle |  |  | Fascicle length |  |  |
| --- | --- | --- | --- | --- | --- | --- | --- | --- | --- |
|  | *ROM,*<br>*LG* | *Jump,*<br>*LG* | *ISO,*<br>*VL* | *ROM,*<br>*LG* | *Jump,*<br>*LG* | *ISO,*<br>*VL* | *ROM,*<br>*LG* | *Jump,*<br>*LG* | *ISO,*<br>*VL* |
| **MAD** | 0.03 cm | 0.03 cm | 0.05 cm | 1.76° | 2.09° | 1.97° | 0.45 cm | 0.58 cm | 0.75 cm |
| **RMSD** | 0.03 cm | 0.04 cm | 0.10 cm | 2.12° | 2.84° | 2.60° | 0.58 cm | 0.73 cm | 1.04 cm |
| **ICC** | 0.95 | 0.93 | 0.97 | 0.89 | 0.87 | 0.82 | 0.94 | 0.89 | 0.76 |

Mean-absolute difference (MAD), root-mean-square difference (RMSD) and intraclass correlation (ICC) between manual estimates from different human observers (n = 3) regarding muscle thickness, pennation angle and fascicle length of lateral gastrocnemius (LG) during range-of-motion (ROM) and jumping, and of vastus lateralis (VL) during isometric contraction (ISO).

**Fig 6.**
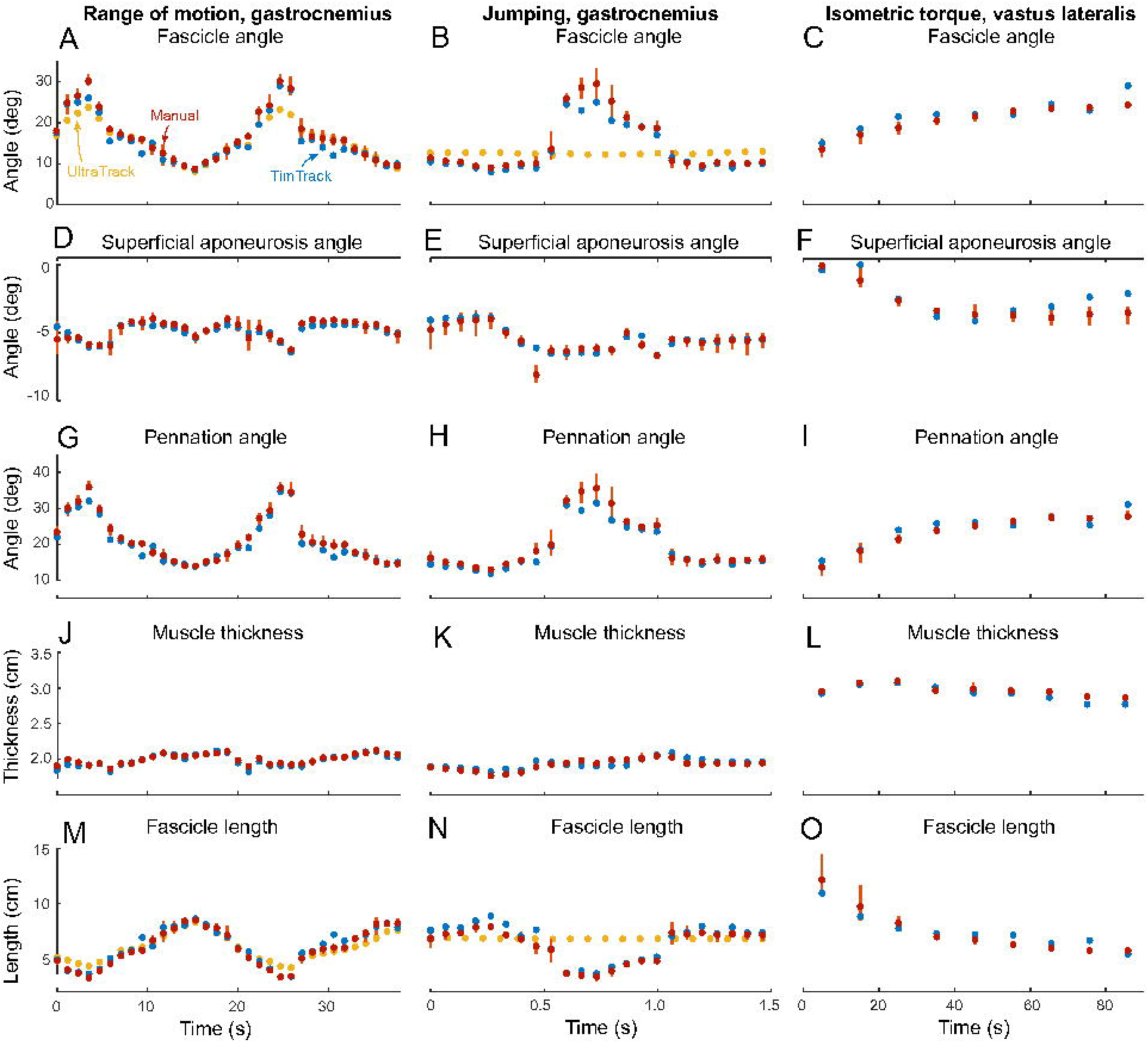
Estimates of TimTrack algorithm and manual observer on geometric muscle features. Estimates of TimTrack algorithm (blue dots) and manual observer (red dots) for fascicle angle (A-C), superficial aponeurosis angle (D-F), pennation angle (G-I), muscle thickness (J-L) and fascicle length (M-O) during range of motion movement (left column), jumping (middle column) and isometric trials (right column). Manual estimates that correspond to the same image are connected by vertical lines. When manual estimates are similar, this connecting line may not be visible. Similarly, when manual and TimTrack estimates are nearly identical, the latter may not be visible.

The TimTrack algorithm’s results were also quantitatively comparable to manual estimates, with differences on the same order as differences between human observers performing manual estimates (RMSE in Table 3 vs. RMSD in Table 2). The algorithm estimate of muscle thickness, pennation angle and fascicle length increased with (the mean of the) observer estimate (slope = 0.99 ± 0.01, 0.99 ± 0.01 and 0.99 ± 0.01 respectively, estimate ± 95% confidence interval, linear regression, see Fig 7). In addition to all regression slopes being close to 1, the algorithm vs. manual estimates of thickness, pennation and fascicle length were well explained by the identity lines (R^2^ = 0.96, R^2^ = 0.84 and R^2^ = 0.83, respectively), indicating that there was negligible systematic under- or overestimation.

**Table 3:**
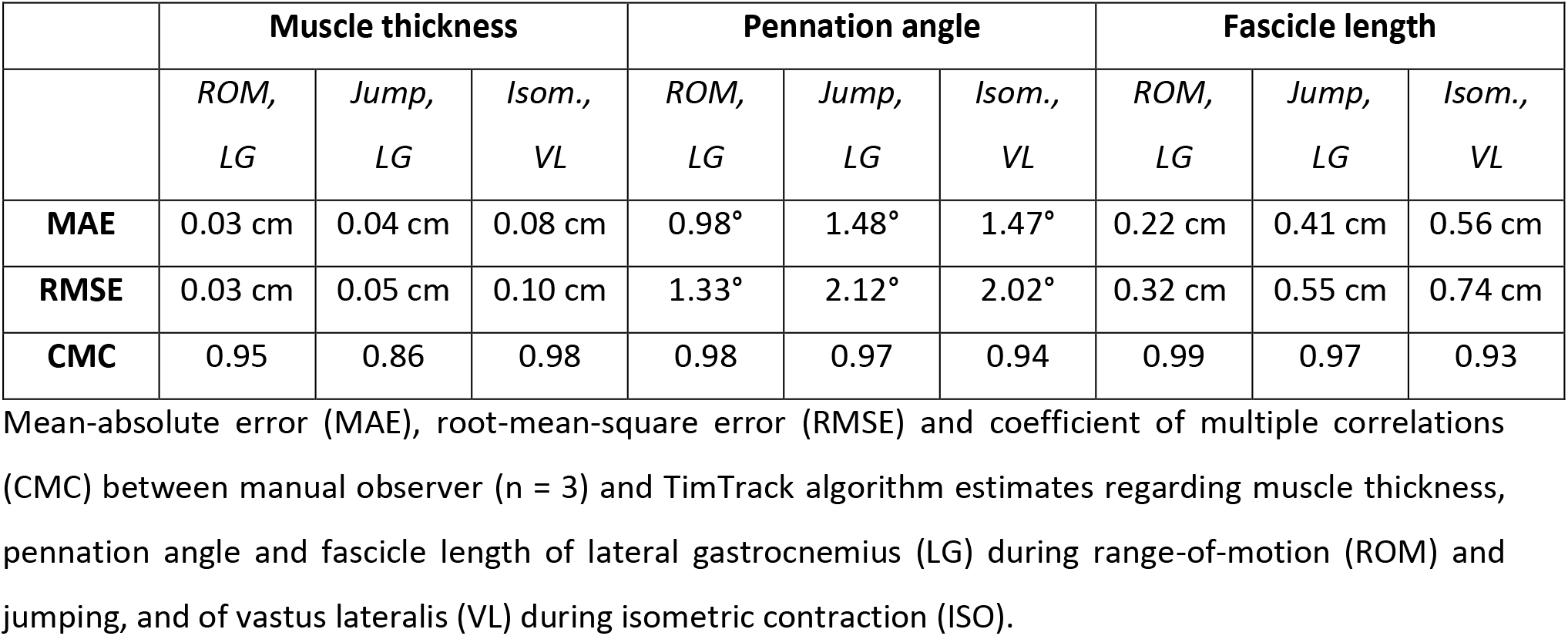
Differences between manual observer and TimTrack algorithm estimates of muscle thickness, pennation angle and fascicle length in ROM, jumping and isometric trials.

|  | Muscle thickness |  |  | Pennation angle |  |  | Fascicle length |  |  |
| --- | --- | --- | --- | --- | --- | --- | --- | --- | --- |
|  | *ROM,*<br>*LG* | *Jump,*<br>*LG* | *Isom.,*<br>*VL* | *ROM,*<br>*LG* | *Jump,*<br>*LG* | *Isom.,*<br>*VL* | *ROM,*<br>*LG* | *Jump,*<br>*LG* | *Isom.,*<br>*VL* |
| **MAE** | 0.03 cm | 0.04 cm | 0.08 cm | 0.98° | 1.48° | 1.47° | 0.22 cm | 0.41 cm | 0.56 cm |
| **RMSE** | 0.03 cm | 0.05 cm | 0.10 cm | 1.33° | 2.12° | 2.02° | 0.32 cm | 0.55 cm | 0.74 cm |
| **CMC** | 0.95 | 0.86 | 0.98 | 0.98 | 0.97 | 0.94 | 0.99 | 0.97 | 0.93 |

Mean-absolute error (MAE), root-mean-square error (RMSE) and coefficient of multiple correlations (CMC) between manual observer (n = 3) and TimTrack algorithm estimates regarding muscle thickness, pennation angle and fascicle length of lateral gastrocnemius (LG) during range-of-motion (ROM) and jumping, and of vastus lateralis (VL) during isometric contraction (ISO).

**Table 4:**
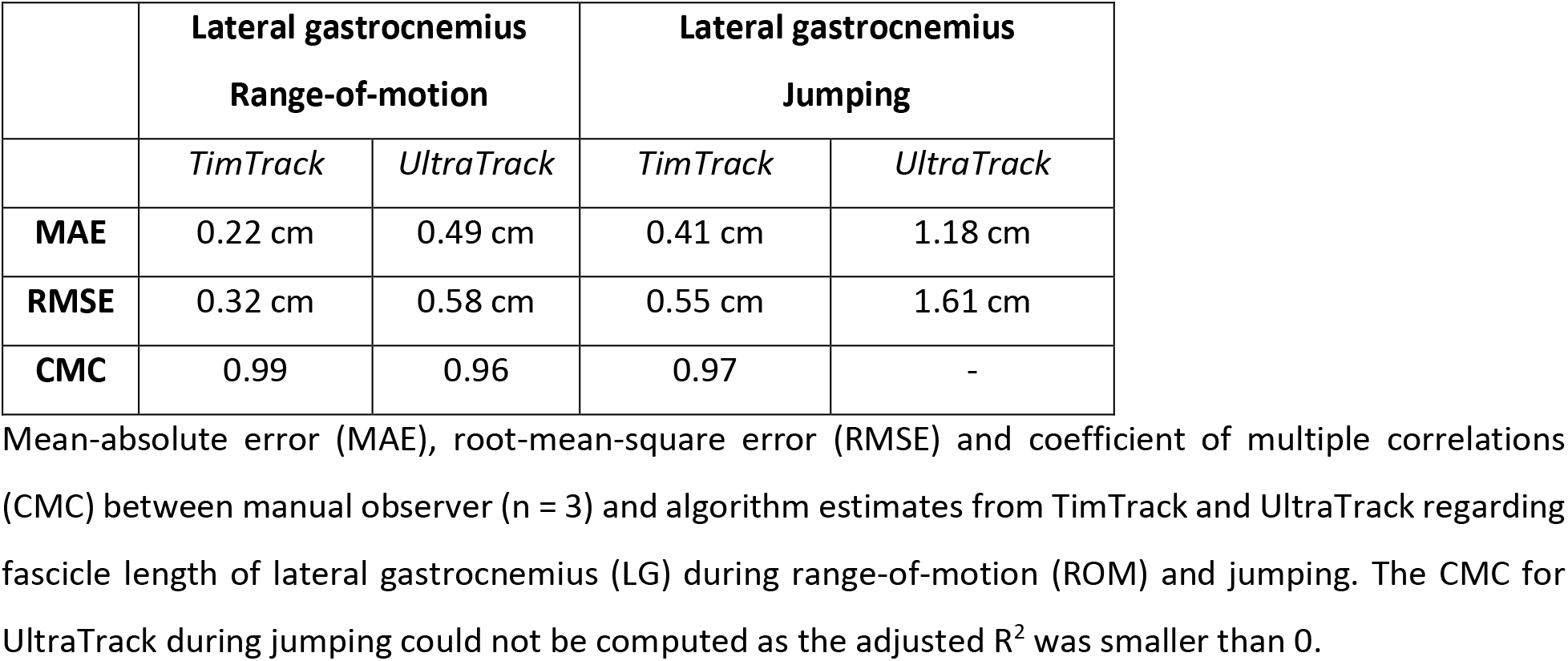
Differences between manual observer and TimTrack algorithm estimates of gastrocnemius fascicle length compared to Ultratrack.

|  | Lateral gastrocnemius<br>Range-of-motion |  | Lateral gastrocnemius<br>Jumping |  |
| --- | --- | --- | --- | --- |
|  | *TimTrack* | *UltraTrack* | *TimTrack* | *UltraTrack* |
| **MAE** | 0.22 cm | 0.49 cm | 0.41 cm | 1.18 cm |
| **RMSE** | 0.32 cm | 0.58 cm | 0.55 cm | 1.61 cm |
| **CMC** | 0.99 | 0.96 | 0.97 | - |

Mean-absolute error (MAE), root-mean-square error (RMSE) and coefficient of multiple correlations (CMC) between manual observer (n = 3) and algorithm estimates from TimTrack and UltraTrack regarding fascicle length of lateral gastrocnemius (LG) during range-of-motion (ROM) and jumping. The CMC for UltraTrack during jumping could not be computed as the adjusted $R^2$ was smaller than 0.

**Fig 7.**
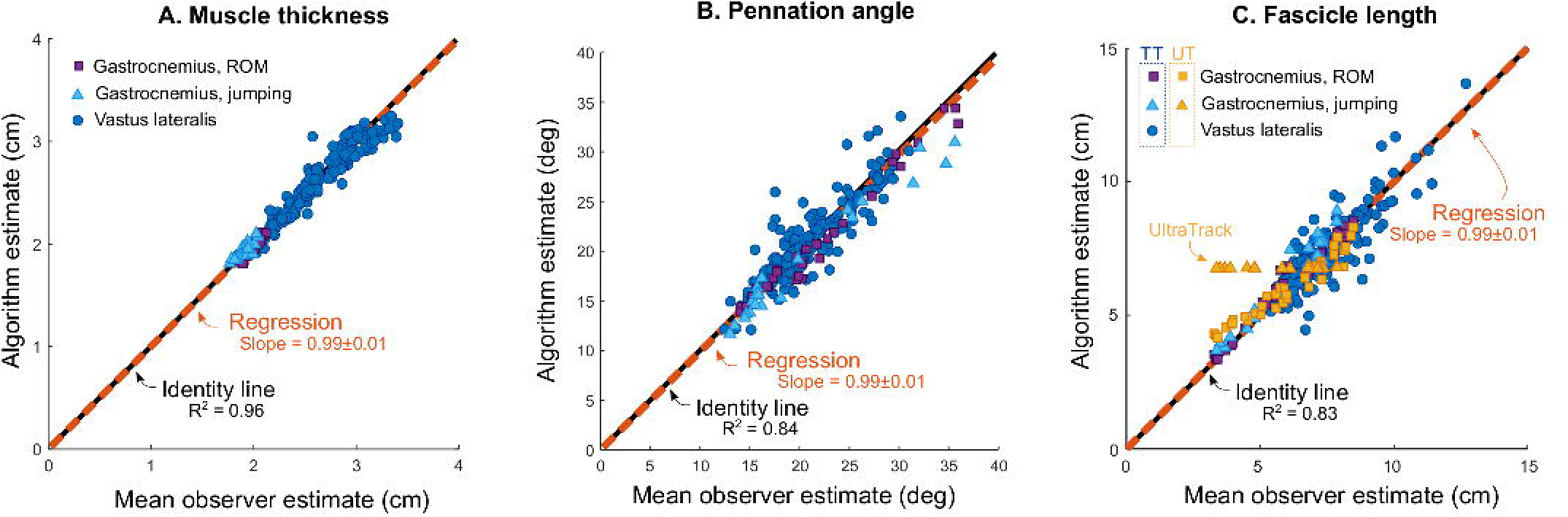
TimTrack algorithm estimates vs. mean manual observer estimates. A. TimTrack algorithm estimates of muscle thickness increased linearly with mean of manual estimates (slope = 0.99±0.01, linear regression). B. TimTrack algorithm estimates of pennation angle increased linearly with mean of manual estimates (slope = 0.98±0.01, linear regression). C. TimTrack algorithm estimates of fascicle length increased linearly with mean of manual estimates (slope = 0.99±0.02, linear regression). TimTrack’s estimates were somewhat more accurate than UltraTrack’s, in particular for gastrocnemius during jumping. Overall, TimTrack estimates were closely related to mean observer estimates (R^2^ = 0.83-0.96).

## Discussion

Our algorithm estimates geometric muscle features from ultrasound images, including muscle thickness, superficial aponeurosis angle, fascicle angle, fascicle length and pennation angle. We found reasonably good agreement between the TimTrack algorithm and manual estimates for human lateral gastrocnemius and vastus lateralis muscle, with algorithm-manual differences comparable to within-manual differences. This indicates that TimTrack may be suitable for replacing manual estimation in experimental studies where these geometric muscle features are of interest. In the following, we compare TimTrack to other contemporary algorithms, considering the respective assumptions, limitations, and performance.

TimTrack provides a high level of automation while combining state-of-the-art features from several other algorithms. We found good accuracy to be achieved with the use of Frangi-type Hessian filtering in combination with Hough transform for fascicle detection [18]. We combined this technique with automatic aponeurosis detection for muscle thickness estimation, adopting an object detection method previously used for fascicle detection [17]. The algorithm by Zhou et al. [20] also includes automated aponeurosis detection, resulting in fascicle length estimates that correlate well with manual estimates (r = 0.92 – 0.98). Their algorithm uses the Radon transform for aponeurosis detection, in contrast to the object detection method used in TimTrack. Using the Radon transform may lead to overestimation of muscle thickness, which has been reported when using the closely-related Hough transform [14]. Furthermore, the algorithm by Zhou and colleagues selects the most dominant fascicle angle from the Hough transform, while TimTrack can uses the median of several dominant angles. We prefer the latter because we found it to yield better results for images with non-uniform fascicle orientations, when the dominant orientation does not necessarily reflect the overall fascicle angle. We did not have opportunity to compare accuracy with the algorithm of Zhou et al. [20], which is not freely available for direct comparison. Altogether, the proposed algorithm combines several approaches to improve objectively, reproducibility and time efficiency.

TimTrack is fundamentally different from optic flow-based algorithms [11–13] as its estimates do not depend on a succession of ultrasound images. The main advantage of optic flow-based algorithms is that they are relatively insensitive to (speckle) noise, and therefore have low requirements on image quality. This is because optic flow only detects global movement, while local (speckle) noise is effectively filtered out. Their main disadvantage is sensitivity to integration drift, which is usually manually corrected for longer image sequences, for example the gastrocnemius lateralis images examined here (see Fig 5). Here TimTrack yields higher accuracy agreeing well with human observers, compared to a state-of-the-art optic flow algorithm (Farris & Lichtwark, 2016). This was accomplished with minimal manual intervention, to set the regions of interest for aponeurosis detection, and without need for manual drift correction or initial fascicle identification. Moreover, TimTrack can optionally extrapolate beyond the image frame if appropriate, as demonstrated on the relatively long fascicles of the vastus lateralis muscle. TimTrack can achieve comparable accuracy to optic flow methods, while having some advantages with respect to drift and manual corrections.

There are nonetheless limitations and sensitivities to TimTrack. For example, image speckle may inadvertently cause a sub-population of fascicles within an image to dominate the calculations, and can thus affect the estimated fascicle orientation from the Hough transform. TimTrack therefore should work best on ultrasound images with relatively high quality and little speckle. Although TimTrack allows for curved aponeuroses, it presently includes only relatively simple, first- or second-order curvature models. It also assumes fascicles to be linear, similar to the other algorithms discussed here. It is therefore challenging to estimate lengths of curved fascicles, especially when there is substantial speckle. Fortunately, muscle fascicle curvature is thought to be most substantial mainly at higher force levels [25]. The application of automated methods, including ours, should therefore be limited to movements where fascicles are fairly linear.

## Conclusion

We here present an automated ultrasound algorithm that provides estimates of geometric muscle features without drift sensitivity of optic flow algorithms, and with more automation than most previous algorithms. Automation allows for relatively fast and objective analysis of many images or image sequences, while retaining accuracy comparable to manual estimations. These features may prove advantageous for cyclic movements such as locomotion, or movements with long image sequences. The TimTrack algorithm estimates muscle geometric features with good accuracy, as tested under force conditions comparable to locomotion.

## Competing interests

The authors declare no competing or financial interests.

## Funding

This work supported in part by the Natural Sciences and Engineering Research Council of Canada (NSERC Discovery and Canada Research Chair, Tier 1) and the Dr. Benno Nigg Research Chair in Biomechanics.

## Data availability

An open-source software implementation of the TimTrack algorithm is freely available. It is implemented in the Matlab software environment (MathWorks, Inc., Natick, MA) and includes typical examples of ultrasound data for demonstration purposes. The TimTrack software is available on: https://github.com/timvanderzee/ultrasound-automated-algorithm

## Notes

https://github.com/timvanderzee/ultrasound-automated-algorithm

### Competing Interest Statement

The authors have declared no competing interest.

### Summary of Updates

Improved vessel enhancement filtering to reduce sensitivity to edge effects. Added option to evaluate geometric muscle features (e.g. muscle thickness) at any specified horizontal location, or using 'extrapolation mode'. Added several figures to clarify the methods, revised all other figures. Included data on lateral gastrocnemius during two movements: jumping and range-of-motion. Included manual estimates from three independent human observers and estimates from competitor ultrasound algorithm for comparison.

