## Supplemental Table 1 for "TimTrack: A drift-free algorithm for estimating geometric muscle features from ultrasound images"

### Supporting information

The parameters and the values used for the current data are listed in Table I. When using the algorithm on similar ultrasound images, most parameter values may be left unchanged. However, for considerably different images, parameter changes may be required. We provide a MATLAB-based software tool to easily change parameters, which is provided in the repository (see Data availability). This tool allows for changing parameters on a typical example image and observing their effect right away. Lateralis gastrocnemius and vastus lateralis ultrasound images were similar enough that the same parameters could be used. The only exception to this was the parameter $o_{\mathrm{deep}}$, set to 1 for vastus lateralis (i.e. linear fit) and to 2 for gastrocnemius (i.e. quadratic fit). This decision was made because gastrocnemius deep aponeurosis was not very straight, whereas vastus lateralis aponeurosis was generally straight.

The region selection parameters $D_{\mathrm{superficial}}$ and $D_{\mathrm{deep}}$ relate to the depth of the image, as some depth settings may cause the aponeuroses to reside outside the upper and lower halves of the image, respectively. The filtering parameters relate to the brightness and resolution of the collected image. For example, line thickness parameters for fascicles and aponeuroses ($\sigma_{\mathrm{fas}}$ and $\sigma_{\mathrm{apo,H}}$) relate to the (average) thickness of these structures in terms of pixels, which depends on image resolution. Threshold parameters for fascicles and aponeuroses ($T_{\mathrm{fas}}$ and $T_{\mathrm{apo}}$) relate to the brightness of these structures, which depends on the overall brightness of the image. Parameters for aponeurosis detection and fitting, and fascicle angle estimation relate to muscle properties and to image quality. For example, some muscles may have aponeurosis and fascicle angles that are outside the range use here ($\beta_{\mathrm{max}}$, $\gamma_{\mathrm{max}}$ and $\theta_{\mathrm{range}}$). The right amount of line angles γ selected in Hough transform ($K$) may depend on image quality, as fewer lines may be available in images with poorer quality.

| Step 0: Region selection | | | |
|---|---|---|---|
| **Parameter** | **Description** | **Value** | **Step** |
| $D_{\mathrm{superficial}}$ | Normalized superficial aponeurosis depth range | 0.05-0.40 | 0 |
| $D_{\mathrm{deep}}$ | Normalized deep aponeurosis depth range | 0.60-0.95 | 0 |
| Step 1: Filtering | | | |
| $\sigma_{\mathrm{fas}}$ | Standard deviation Gaussian Hessian fascicles | 1-2 pixels | 1.1a |

| | | | |
|---|---|---|---|
| $\sigma_{\mathrm{apo,H}}$ | Standard deviation Gaussian Hessian aponeurosis | 18-20 pixels | 1.1b |
| $T_{\mathrm{fas}}$ | Threshold for fascicles | 50% brightest | 1.2a |
| $\sigma_{\mathrm{apo,k}}$ | Standard deviation Gaussian kernel aponeurosis | 10 pixels | 1.3 |
| $T_{\mathrm{apo}}$ | Threshold for aponeurosis | 50% brightest | 1.1c & 1.4 |
| $L_{\mathrm{ratio,max}}$ | Maximum length ratio between longest and second-longest aponeurosis objects | 0.9 | 1.4 |
| Step 2: Aponeurosis point detection and fitting | | | |
| $x_{\mathrm{margin}}$ | Margin of horizontal aponeurosis point | 20 pixels | 2 |
| $n_{\mathrm{apox}}$ | Number of points in aponeurosis points | 10 | 2 |
| $o_{\mathrm{deep}}$ | Order of the deep aponeurosis fit | 1 or 2 | 2 |
| $\beta_{\mathrm{max}}$ | Maximal angle of superficial aponeurosis fit | 0.5° | 2 |
| $\gamma_{\mathrm{max}}$ | Maximal angle of deep aponeurosis fit | 0.5° | 2 |
| Step 3: Fascicle angle estimation | | | |
| $w_{\mathrm{ratio}}$ | Ratio between ellipsoid width and image width | 0.80 | 3.1 |
| $\theta_{\mathrm{range}}$ | Range of Hough angles $\theta$ considered | 55-80° | 3.2 |
| $\theta_{\mathrm{res}}$ | Angle resolution in Hough transform | 1° | 3.2 |
| $K$ | Amount of selected line angles $\gamma$ | 10 | 3.4 |

**Table I: Parameter values**
